# An atlas of natural killer cell receptor expression on healthy donor T and NK lymphocytes

**DOI:** 10.1101/2025.09.21.677654

**Authors:** Stacey N. Lee, Derek Rowter, Jeanette E. Boudreau

## Abstract

Human T cells and natural killer (NK) cells exhibit variegated co-expression of germline-encoded receptors, which diversifies their effector functions. CD8+ T cell expression of receptors more typically associated with NK cells has been noted, but a systematic analysis of their distribution has not been described. Here, we comprehensively measured human NK cells and T cells’ expression of NK cell-associated receptors to define their co-expression and patterns associated with donor sex, maturation, and activation.

To assess the activating and inhibitory receptor repertoire of human T cells and NK cells, we developed a 25-colour flow cytometry panel that included channels based on receptor functions. Since NK cell function is known to be driven by interactions with HLA supergroups (KIR ligands), we stratified donors based on KIR-HLA allelic combinations. NK and CD8+ T cell phenotypes and responsiveness were assessed at rest, and in response to the missing self target cell line, K562. We find that NK cells universally express CD45RA, CD161, and the inhibitory receptors TIGIT/TIM3/LAG3. On CD56^dim^CD16^high^ NK cells (those most aligned with missing self reactivity, CD161, CD45RA, natural cytotoxicity receptors (NCRs), and TIGIT/TIM3/LAG3 were most frequently expressed. Not all educated NK cells respond to missing self-targets; those that did exhibited high expression of NCRs, NKG2C, IL-7R/IL-18Ra, and TIGIT/TIM3/LAG3, and lower expression of DNAM-1 and CCR7, in addition to the KIR molecules that defined their status as educated. We note that up to 50% of NK cells express CD8, and this population co-expressed CD16, NCRs, and KIR, and exhibited greater cytotoxicity than the CD8-NK cell population. Among CD8^+^ T cells, acquisition of NK cell-associated receptors was increased as they progressed through differentiation states: effector memory and terminally-differentiated CD8^+^ T cells exhibiting higher expression of NKG2A, KIR, and CD16.

Taken together, the variability of receptor expression patterns highlights the diversity of lymphocyte populations and suggests shared features among the cytotoxic lymphocytes.

## INTRODUCTION

To combat intracellular infections, recognize damaged or transformed cells, and maintain immune homeostasis, human lymphocytes are endowed with collections of receptors that convey signals for activation, inhibition and mobility. Typically, the two major cytotoxic lymphocyte populations – natural killer (NK) and CD8^+^ T cells – are considered as co-operative, but independent in their function and receptor use. NK cells, innate-like lymphocytes, recognize target cells through the binding of germline encoded activating and inhibitory ligands. NK cells are particularly known for their diverse constellation of receptors, which informs how a given NK cell will respond to a challenge, and a major function to recognize target cells with aberrant expression of “self” Human Leukocyte Antigens (HLA) (1, 2). CD8^+^ T cells instead employ antigen-specific T cell receptors (TCRs), whose germline-rearranged variable regions depend on presentation of specific antigen in self HLA molecules. Recent reports demonstrate that some subsets of T cells express receptors canonically exhibited on NK cells, but how these receptors reflect patterns in NK cells or whether they contribute to CD8+ T cell function is unknown.

Permutations of receptor expression enable high phenotypic diversity among NK cells, with as many as 30,000 different subsets present in each individual (2). NK cells are grossly classified by their expression of CD56 and the Fc receptor, CD16, but modern techniques enable their further division into distinct functional subsets, including tissue resident, adaptive, and immunoregulatory, with functions shaped by the receptors they express (3–6). Killer-immunoglobin like receptors (KIR) are expressed only on a subset of NK cells and encoded separately for the genes encoding their HLA I ligands. Co-expression of a KIR allele and its cognate ligand (supergroups of HLA I molecules termed “KIR ligands”) enables missing self reactivity and these cells are termed “educated” (7). NK cells found within lymphoid tissues express higher levels of the chemokine receptor CCR7, while NK cells expressing NKG2C may have transitioned to a more adaptive phenotype (6, 8, 9). Although expression of NK cell receptors is fluid and can be altered in response to the cytokine exposure, triggering of activation pathways, or interactions with specific pathogens, but is largely conserved over an individual’s adulthood (6, 10–12).

Although NK cells are lymphocytes, they are not typically considered among the “adaptive” ones, whose most prominent receptors are germline-rearranged and highly antigen-specific. Both CD4+ “helper” and CD8+ “cytotoxic” T cells rely on the presentation of antigen in “self” HLA I, which activates their T cell receptor (TCR) (13). CD4^+^ T cells, upon activation and in response to environmental cues, are directed into specialized subsets with major roles in priming and polarizing immunity (14). CD8^+^ T cells drive immunity against intracellular pathogens, and exhibit overlapping functions with NK cells: similar cytokine production and cytotoxicity, with both cell types executing target cell killing through the release of perforin and granzyme B (15, 16). Both CD4^+^ and CD8^+^ T cells can differentiate into memory cells, with further priming for tissue residency, circulation and renewal (17). T cells’ repertoires of receptors are not known to be as diverse as those of NK cells, but they nevertheless share inhibitory receptors including the checkpoint receptors PD-1 and TIGIT, and cytokine receptors that mark them as distinct subsets (18–20).

Cytotoxic CD8^+^ T cells and NK cells are often considered as two sides of an innate-adaptive coin: CD8^+^ T cells rely on HLA I presentation; downregulation of HLA I creates a missing self target for NK cells. In this way, the cytotoxic subsets perform complementary functions to force expression of HLA (and antigen), or eliminate cells in which it may be compromised. It is becoming increasingly evident, however, that CD8^+^ T cells can use receptors typically expressed on NK cells, and that NK cells can acquire expression of CD8^+^ as they gain cytotoxic function (21–24). These observations imply that a greater overlap among the cytotoxic subsets might be present. To begin to explore these considerations, we used high parameter flow cytometry to examine the receptor repertoire of circulating CD4^+^ and CD8^+^ T cells and NK cells from healthy adult human donors, focusing on receptors that are canonically associated with NK cells.

NK cell receptor expression is consistent regardless of age or sex but CD4^+^ T cell frequency increases, while CD8+ frequency decreases with age (but not sex). NK cells bearing receptors typically associated with maturation (i.e. KIR, NKG2C, and CD16) exhibited increased expression of CD8, and greater degranulation (CD107a) against the missing self target K562 compared with their CD8-negative counterparts. CD8^+^ T cells acquired expression of several NK cell-associated receptors as they enter a more terminally differentiated state, with effector memory and terminally differentiated effector memory T cell re-expressing CD45RA cell (TEMRA) subsets expressing NKG2A, KIR, and CD16. Taken together, these results demonstrate a shared use of receptors canonically-associated with NK cells among cytotoxic lymphocytes.

## MATERIALS AND METHODS

### PBMC isolation

Peripheral blood mononuclear cells (PMBC) were isolated from buffy coats collected from healthy human donors via the Canadian Blood Services Blood4Research program. Use of PBMC was approved by the Dalhousie University REB (#2016-3842) and the Canadian Blood Services REB (#2016-016). Human PBMCs were collected from buffy coats using Ficoll gradient centrifugation and stored as previously described (25).

### K562 co-culture to test missing self responsiveness

Isolated human PBMCs were quickly thawed, washed, and rested overnight at 37°C in 5% CO_2_ in RPMI with 100 IU/mL human IL-2 (Peprotech) at 1x10^6^ cells/mL to allow for cell recovery. The HLA-negative target cell line K562 (American Type Culture Collection (ATCC)) was cultured at 37°C in 5% CO_2_ in RPMI-1640 media with 10% heat inactivated FBS, 100 U/ml penicillin, and 100 mg/ml streptomycin. In a 96-well v-bottom plate, resuspended PBMCs was added to K562 (effector:target ratio of 3:1) and mixed with gentle pipetting to ensure proper resuspend of the culture. They were the co-culture in the presence of anti-LAMP1 (CD107a, H4A3, BD Biosciences) for 5 h at 37°C. The plate was thereafter centrifuged at 300 *g* x 5 min, rinsed with 200 μL PBS, and stained for flow cytometry.

### Flow cytometry

Our flow cytometry panel was validated through the use of known positive and negative expressing cells for each antibody target. All antibodies were titrated to optimal dilution for each panel and instrument and confirmed to not infer with fluorescence in other channels through fluorescence minus one (FMO) controls. Flow cytometry surface staining was conducted in a V-bottom 96-well plate, and each incubation was done in the dark at RT. Following each washing step, the cells were centrifuged at 300 x *g* for 5 min at RT. All samples, except the unstained control, were resuspended in 50 μL of fixable viability dye and incubated for 20 min. Antibody cocktails (Supplementary table 1) were then prepared either just fluorescence-activated cell sorting wash buffer ((FWB) (PBS, 1% bovine serum albumin (ThermoFisher), 0.1% sodium azide (Sigma Aldrich)) and 10 μL per well (50 μL) of Brilliant Stain Buffer Plus (BD Biosciences). Following viability staining, 150 μL of PBS was added to each well, and the plate centrifuged (300 *g* x 5 min). To mitigate non-specific antibody binding, samples were resuspended in Fc Block (1/50 dilution in PBS, BD Biosciences) and incubated for 20 min.

Following this incubation, 150 μL of PBS was added and centrifuged as described. Samples - excluding unstained controls, viability controls, or other stained controls - were resuspended in 50 μL of the staining cocktail and incubated for 30 min. Following this incubation, 150 μL of FWB was added to each well, centrifuged, and the samples were washed once more with 200 μL of FWB. Cells were then fixed with 100 μL of 4% paraformaldehyde for 10 min, then washed twice with 200 μL of FWB and resuspended for flow cytometry sample acquisition.

FACS acquisition was performed using a BD FACS Symphony analyzer or BD FACS Fortessa analyzer. To normalize acquisition metrics between experiments, 8-peak Rainbow Sphero beads (BD Biosciences) were used and laser voltages adjusted to ensure equivalent mean fluorescence intensities (MFI) across experiments. All exported files were compensated and analyzed using FlowJo v.10.10.1 (BD Biosciences). Uniform Manifold Approximation and Projection (UMAP)s were made in FlowJo, by first downsampling all samples to the same cell number, followed by concatenation of the downsampled populations. A UMAP of the concatenated output was then made using the FlowJo plugin with default settings.

Regardless of cell source, all samples were initially gated on forward scatter area (FSC-A) vs side scatter area (SSC-A) within the area that the target cells were expected. Doublets were excluded first by gating on FSC-height (FSC-H) by FSC-width (FSC-W), and gating on SSC-height (SSC-H) by SSC-width (SSC-W). Finally, viable cells were identified by gating on viability dye by FSC-A. True positive gating of all markers of interest was conducted by using the viability control as the true negative population and with gate drawn to the right of the unstained viable population. Viable NK cells were identified as CD56+CD3-with subsequent gating to identify key subpopulations. A representative gating scheme for gating on lymphocytes (Supplementary figure 1) is shown below.

### Statistical analysis

Expression data measured by flow cytometry show the mean ± the standard deviation with significance indicated in the figure, unless otherwise specified. All statistical computations were performed using GraphPad Prism [Version 9]. Statistical significance was set at p<0.05, and significance levels are defined in figures as *, p<0.05; **, p<0.01, ***, p<0.001; ****, p<0.0001. Statistical tests and significance are indicated within individual figure captions.

Normal distribution was assumed in experiments with eight donors or more. When comparing differences between two groups, nonparametric paired *t* tests were used. In experiments comparing variation between >2 parameters with one independent variable, one-way analysis of variance (ANOVA) was completed with multiple comparisons. When comparing variation between >2 parameters with two independent variables ordinary two-way ANOVA with multiple comparisons was used. In the case where there was an uneven number of samples (such as when separating donors by sex), mixed-effects analysis was used with multiple comparisons.

## RESULTS

### NK cell populations are consistent across sex and age

The receptors expressed by a given lymphocyte dictate its function, level of maturation, and activation status. Using a 25-colour high parameter flow cytometry staining panel that marked 35 receptors, we examined 30 healthy donors peripheral NK and T cell repertoires (Supplementary Table 2.1). T cells represented majority of circulating lymphocytes (80.12%±8.48%), with a higher proportion of CD4^+^ cells (66.07%±11.17%) compared to CD8^+^ cells (29.54%±9.98%) (Figure 1A). NK cells accounted for a lower proportion (7.97%±4.90%) of circulating lymphocytes in this cohort. There was no significant variability associated within the sex of the donors (Figure 1B). When stratified by age, there were significantly more CD4^+^ (71.2%±9.1% vs 61.6%±11.1%) and significantly fewer CD8^+^ T cells (25%±8.2% vs 33.5%±9.9%) in the older cohort of donors (50-72 years) when compared to the younger cohort (22-49 years; Figure 3.1C). We next assessed how age and sex correlated with the expression of 20 receptor groups that included markers of differentiation, activation, and inhibition. Within the CD4^+^ T cell compartment, expression of the cytokine receptors IL7R/IL18Rα (dumpgated) was higher in female donors (55.9%±14.2% vs 48.2%±10.9%) and in the younger cohort (55%±11% vs 47.6%±13.25%; Figure 1D and E). Conversely, CCR7 expression was higher in the older cohort (77.2%±6.4% vs 68.8%±10.25). There was an elevated expression of the combined checkpoint receptor group that included TIGIT_TIM3_LAG3 on CD8+ T cells among female donors (37.6%±15.5% vs 30.8%±11.5%), and decreased expression of CD45RA (66.1%±8.3% vs 53.7%±14.4%) and an increased expression of DNAM-1 (55.2%±10.8% vs 42.2%±13.2%) in the older donor cohort (Figure 1F and G). There were not statistically-significant differences in receptor expression across age or sex for the NK cell subset (Figure 1H and I). Taken together, these findings confirm the known impacts of aging on T cell distribution and that NK cell repertoires are more stable with aging (12).

**Figure 1.**
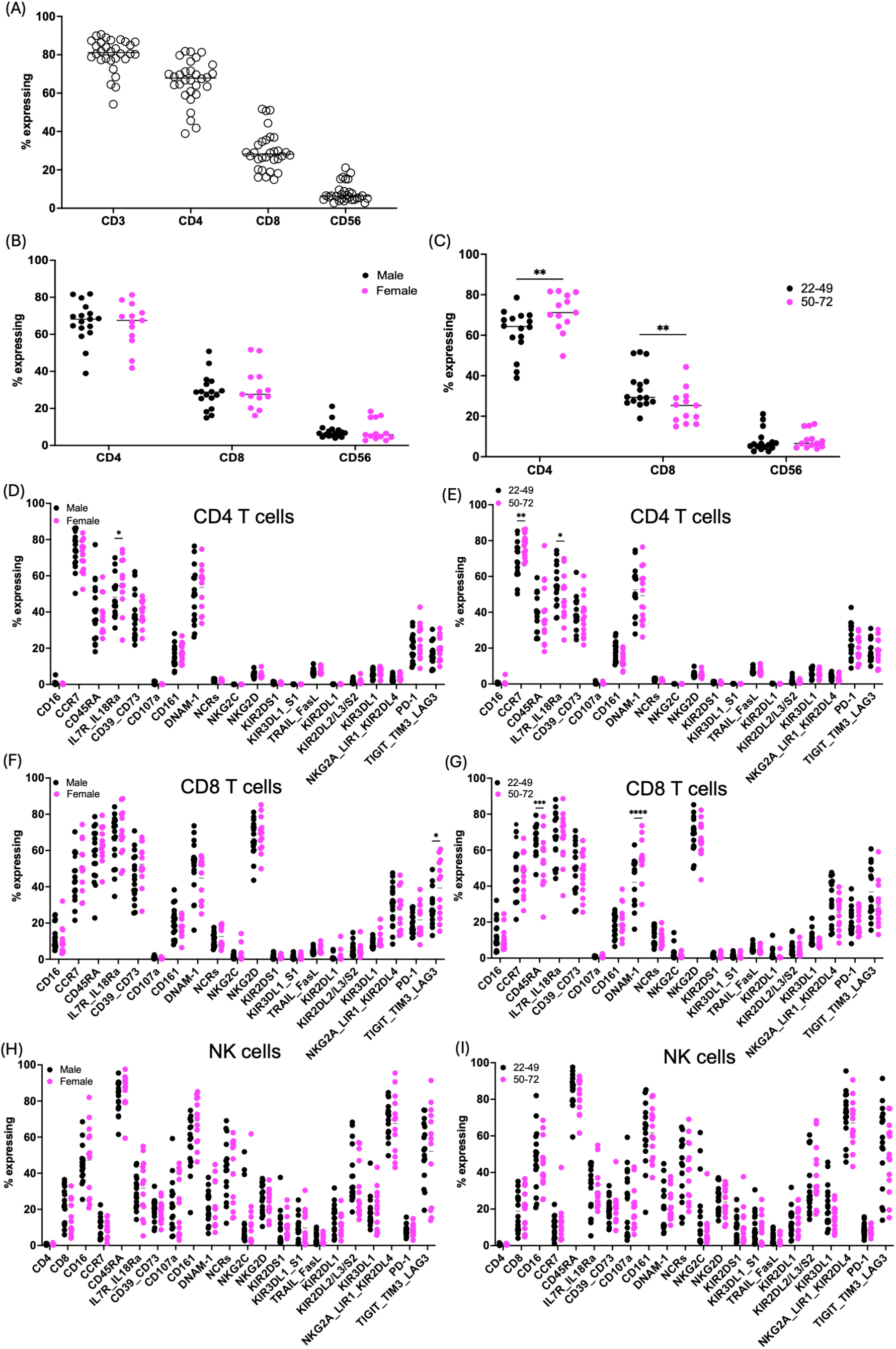
Distribution of the immune cell subtype by donor demographics. Using 25-colour flow cytometry, the immune cell phenotype of healthy human PBMCs was assessed. (A) percentage of viable lymphocytes. (B) percentage of viable lymphocytes stratified by sex. (C) percentage of viable lymphocytes stratified by age. (D-I) Lymphocytes were then further subdivided and the receptor phenotype of each subtype assessed by age (CD4^+^ (D), CD8^+^ (F), and NK cells (H)) and sex (CD4^+^ (E), CD8^+^ (G) and NK cells (I)) (n=30); statistics shown are mixed-effects analysis with multiple comparisons; *=p<0.05, **=p<0.01, ***=p<0.001, ****=p<0.0001

### CD8^+^ T cells express canonically NK cell receptors

NK cells integrate signaling from multiple receptors to determine the ultimate outcome of interactions with target cells (1). To visualize how receptors are co-expressed among lymphocyte subsets, we performed Uniform Manifold Approximation and Projection (UMAP) analysis. To ensure that only the T cell and NK cell populations were considered, CD3^-^CD56^-^ cells were excluded. Using an unsupervised analysis, NK, CD8^+^ and CD4^+^ T cells clustered separately, as expected (Figure 2A). Overlaying each receptors onto our UMAP, we could visualize receptor expression among these subsets, which we quantified within donors (Figure 1B-E). Within the CD4^+^ T cell cluster, CD45RA expression is segregated from the expression of checkpoint receptors PD-1 and TIGIT_TIM3_LAG3, indicating a separation of naïve from terminally differentiated/exhausted populations. This separation of naïve subsets from effector populations is also noted in the CD8^+^ T cell group, as a portion of CCR7 segregates from the checkpoint receptors. Typically NK cell-associated receptors, including killer-immunoglobulin like receptors (KIR), NKG2A and NKG2D, and the Fc receptor CD16, are found within both the NK and CD8^+^ T cell clusters.

**Figure 2.**
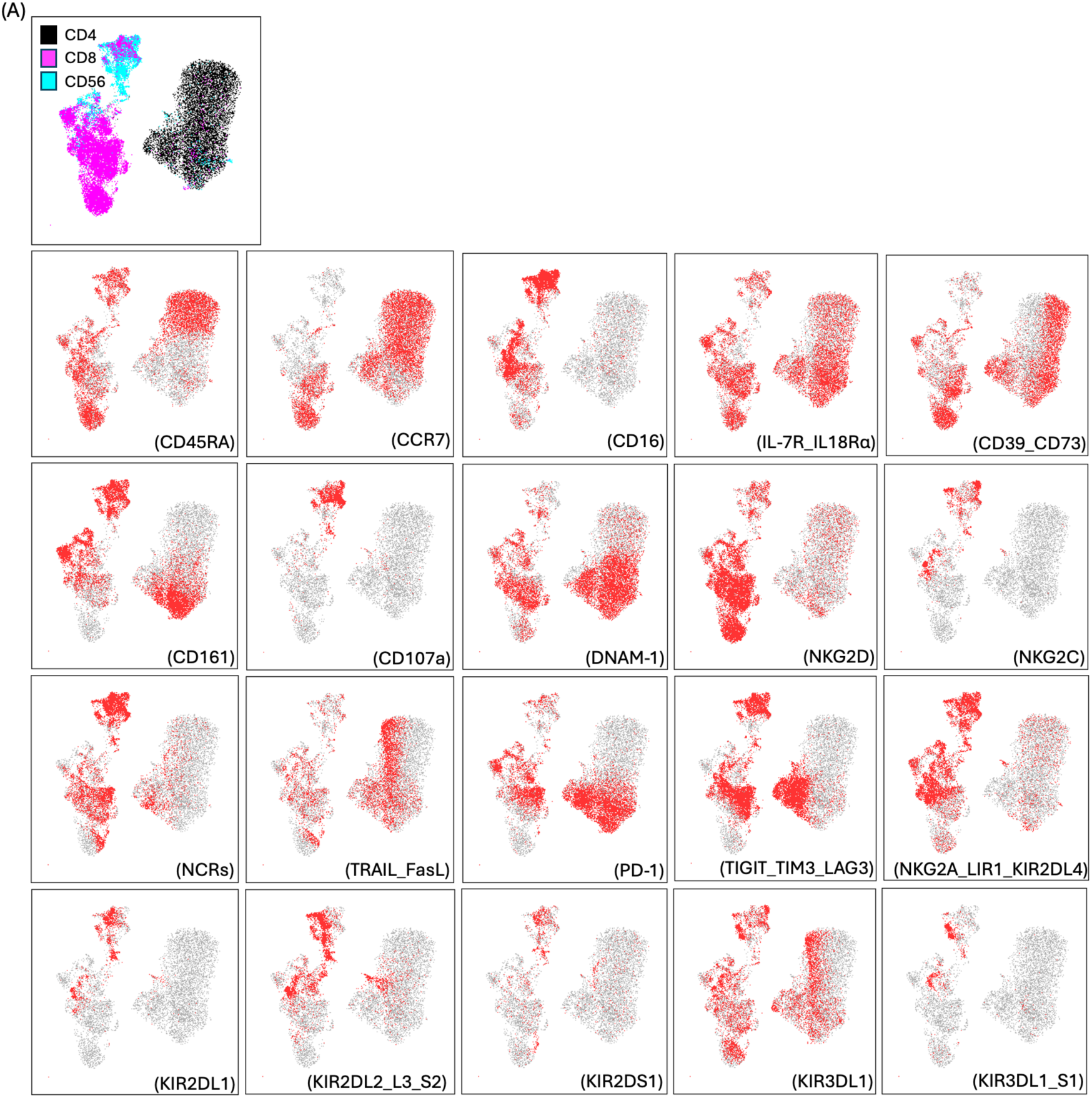
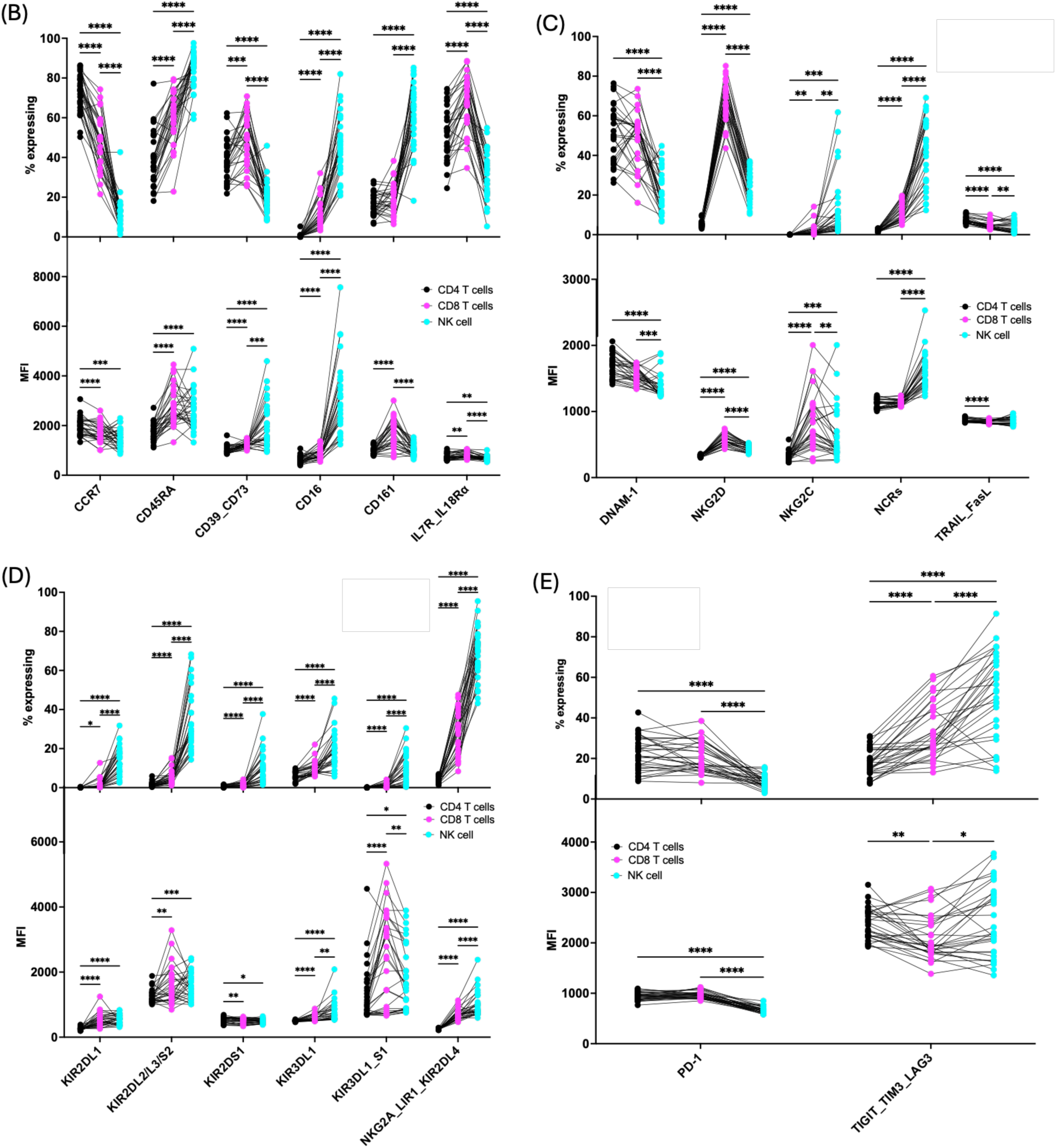
Distribution of receptor expression among T cell and NK subsets. A comparison of the expression of various receptors on CD4^+^ and CD8^+^ T cells and NK cells. (A) UMAP analysis of the lymphocyte population showing distinct T cell and NK cell populations. UMAP overlays demonstrate where the expression of each receptor (red shading) falls within the populations. (B-D) The percent expression (top graph) and MFI (bottom) of each receptor group. Lines connect donors across cell types (B) Differential receptors. (C) Checkpoint receptors. (D) Activating receptors. (D) KIR expression. (n=30); statistics shown are a two-way ANOVA with multiple comparisons; *=p<0.05; **=p<0.01; ***= p < 0.001; ****=p<0.0001

**Figure 3.**
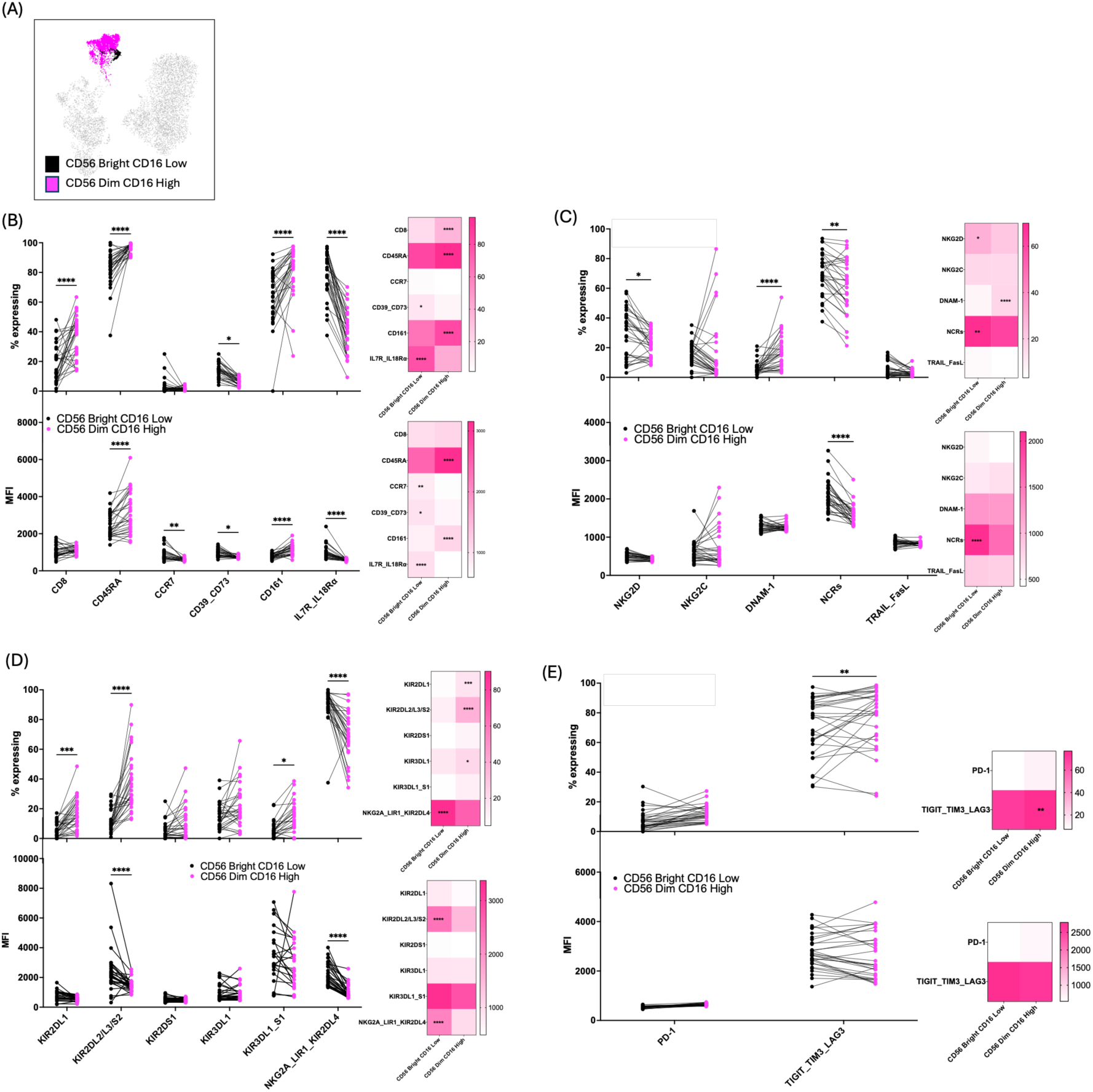
CD56^dim^ CD16^high^ NK cells have increased KIR expression and increased expression of CD8. A comparison of receptor expression between CD56^bright^ CD16^low^ and CD56^dim^ CD16^high^ NK cells. (A) Overlay of CD56^bright^ CD16^low^ and CD56^dim^ CD16^high^ populations of UMAP on the total lymphocyte population. (B-D) The percent expression (top graph) and MFI (bottom) of each receptor group. Lines connect same donors (B) Differential receptors. (C) Checkpoint receptors. (D) Activating receptors. (D) KIR expression. (n=30); statistics shown are a two-way ANOVA with multiple comparisons; *=p<0.05; **=p<0.01; ***= p <0.001; ****=p<0.0001

Immune checkpoint expression was present on both T and NK cells but varied between them. Both T cell subsets expressed more PD-1 than NK cells (CD4^+^: 22.3%±8.5%; CD8^+^: 20.5%±7.3%; NK cells: 8.4%±2.9%), while NK cells expressed a higher proportion of TIGIT/TIM3/LAG3 (CD4^+^: 18.1%±6.5%; CD8^+^: 33.2%±13.5%; NK cells: 52%±21.2%; Figure 1E). Though the adenosine receptors CD39/CD73 were proportionately highest on CD8^+^ T cells, (48.2%±12.9%), their MFI was highest on NK cells, indicating that they are expressed at a higher surface density. Conversely, CD161, a marker frequently expressed on NK cells (61.2%±15.5%), exhibited its highest intensity on CD8^+^ T cells.

NK cells had the highest expression of the activating natural cytotoxicity receptors (NCRs) among lymphocyte populations (NK: 41.05%±16.78%; CD8^+^ T cells: 11.54%±4.56%; CD4^+^ T cells: 2.12%±0.64) (Figure 2C). Both T cell subsets expressed DNAM-1 and CD8^+^ T cells also expressed more NKG2D than NK cells. Finally, though NK cells had a higher expression of KIR (13.8-35.1%±7.1-16%) and CD16 (45.7%±15.1%), CD8^+^ T cells also expressed both of these inhibitory receptors at low levels (KIR: 1-9.7%±1-3.5%; CD16: 11.8%±7.3) (Figure 2B and D). Taken together, these findings reveal typically NK cell-associated receptors as putative features that may influence lymphocyte subset diversification.

### CD56^dim^CD16^high^ NK cells display a more mature phenotype with increased expression of CD8 and KIR receptors

NK cells are typically classified by their surface expression of CD56 and CD16: CD56^bright^CD16^low^ NK cells exhibit cytokine production and tissue residency, while the highly cytotoxic CD56^dim^CD16^high^ constitute the majority of circulating NK cells and considered to be the more mature NK cell subset in the blood (26, 27). Within our dataset, the CD56^bright^CD16^low^ population formed a distinct subcluster among the NK cell cluster on the lymphocyte UMAP (Figure 3A).

Comparing the two major NK cell subsets, we observed that CD56^dim^CD16^high^ subset displayed features of mature NK cells. TIGIT/TIM3/LAG3, which are typically expressed by mature or activated NK cells, was upregulated in the CD56^dim^CD16^high^ population (77%±20.9%; Figure 3E) (28, 29). CD161, which has previously been shown to be expressed on mature NK cells and co-expressed with checkpoint receptors, exhibited both an increased expression (82%±16.4%) and MFI (Figure 3B) (30). Conversely, expression of the activating receptors NKG2D (21.8%±8.7%) and the NCRs (61.3%±18.6%) were decreased in the CD56^dim^CD16^high^ subset (Figure 3C). Notably, CD56^dim^CD16^high^ NK cells displayed increased expression of CD8 (36.2%±14.5%), a marker that has more recently been shown to be associated with a more mature NK cell phenotype (31).

Both expression (78.4%±15.5%) and MFI of IL7R/IL18Rα were higher in the CD56^bright^CD16^low^ population, which echoes previous work demonstrating that IL-7R is predominantly expressed on CD56^bright^ NK cells and that, further, CD56^bright^ NK cells are more responsive to exposure to IL-18 (Figure 3B) (32, 33). Though there was no significant decrease in the percent positive expression, there was a decrease in the MFI of CCR7 on the CD56^dim^CD16^high^ subset, reflecting a loss of CCR7 expression as CD56^bright^ NK cells mature and shift toward a circulating CD56^dim^ phenotype (9, 34).

As the more mature NK cell population, CD56^dim^CD16^high^ have also been associated with increased KIR expression and NK education (35, 36). NK cells distinguish self from non-self through the binding of KIR to HLA I, which is constitutively expressed on all healthy cells (37). Upon binding of their cognate HLA I, these KIR expressing NK cells are endowed with a lower threshold of activation and increased cytotoxicity. Within our cohort, CD56^dim^CD16^high^ NK cells displayed an increased expression of KIR2DL1 (16.8%±10.6%), KIR2DL2/L3/S2 (37.6%±19.3%), and KIR3DL1 (22%±14.4%, p=0.0533; Figure 3D). NKG2A, which is associated with less mature NK cells, was elevated (90.3%±11.2%) in the CD56^bright^CD16^low^ population and decreased (68.9%±16.1%) in the CD56^dim^CD16^high^ population, as expected. The activating receptor DNAM-1, which improves binding and activation of educated NK cells, is further associated with NK cell maturation and was more highly expressed (40%±15.1%) in the CD56^dim^CD16^high^ population (38, 39).

### Missing self responsive NK cells have increased expression of the activating NCRs and checkpoint receptors

The receptor repertoire of a given NK cell can determine whether and how strongly it responds to a target cell. To assess the phenotype of responding NK cells, PBMCs were co-cultured with an HLA class I negative cell line, K562, and stained for degranulation by anti-CD107a (LAMP-1). CD107a^+^ NK cells were relatively diffuse throughout the NK cell cluster (Figure 4A).

**Figure 4.**
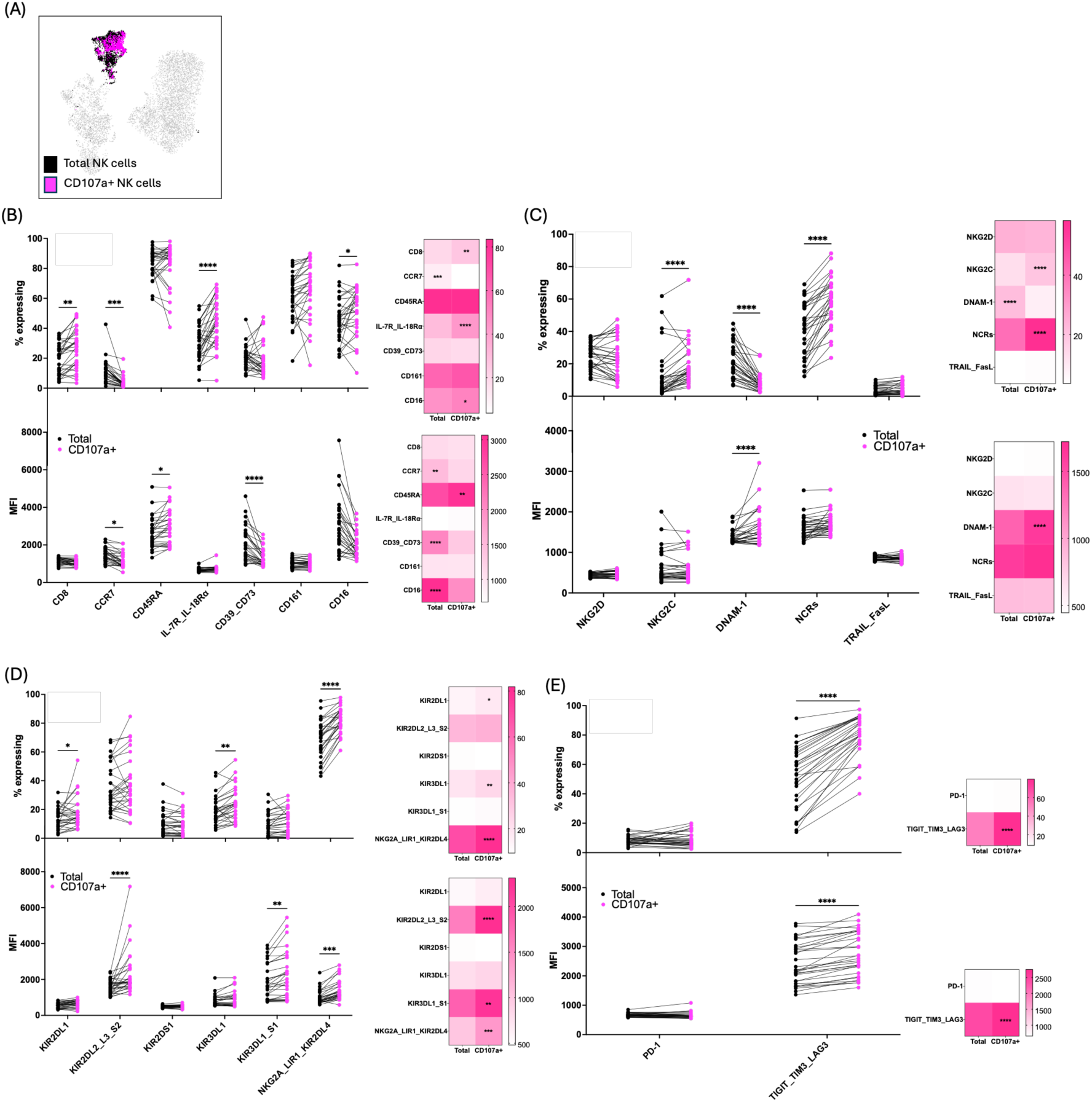
Responding NK cells have increased expression of NCRs and decreased DNAM-1. A comparison of receptor expression between total and responding (CD107a+) NK cell populations. NK cells responsiveness was assessed by cocultured with the HLA negative cell line K562 (3:1 E:T) (A) Overlay of total and responding populations of UMAP on the total lymphocyte population. (B-D) The percent expression (top graph) and MFI (bottom) of each receptor group. Lines connect same donors (B) Differential receptors. (C) Checkpoint receptors. (D) Activating receptors. (D) KIR expression. (n=30); Statistics shown are a two-way ANOVA with multiple comparisons; *=p<0.05; **=p<0.01; ***= p < 0.001; ****=p<0.0001

Responding NK cells were characterized by an increased expression of the NCRs (58.3%±15.6%) when compared to the total NK cell population (41.05%±16.78%; Figure 4C), and higher expression of NKG2C (19.4%±13.6%). Further, CD8^+^ expression was elevated in the responding population (26.3%±13%; Figure 4B).

DNAM-1 was expressed on fewer NK cells (8.1%±5.6%; total: 21.4%±10.6%), but responding NK cells had a higher intensity of expression, as determined by MFI, suggesting a greater density of receptors on each individual cell. The inhibitory receptor TIGIT negatively regulates the activation of DNAM-1 through competitive binding of their shared ligands and through *cis* interactions with DNAM-1 (18, 28). Within our dataset, TIGIT/TIM3/LAG3 was increased in both expression (79.8%±14.1%) and MFI in the responding NK cell subset, potentially indicating that these receptors may be expressed in opposition to one another in the context of activation and that responding NK cells may be shifting from a more activated phenotype to one of exhaustion (Figure 3.4 E) (11).

As an HLA class I negative cell line, K562s are frequently used to test missing-self reactivity of NK cells. Within our responding NK cell population, there was increased expression of KIR2DL1 (17.6%±10.7%), KIR3DL1 (23.8%±11.5%), and NKG2A_LIR1_KIR2DL4 (81.8%±8.9%; Figure 4D). Further, there was an increase in MFI for KIR2DL2/L3/S2, KIR3DL1/S1, and NKG2A/LIR1/KIR2DL4, indicating that within our dataset the KIR^+^ NK cells play an important role in responding to an HLA negative target.

### CD8^+^ NK cells display a more mature phenotype with increased activation

CD8 is a well-defined co-receptor for the TCR of effector T cells, but emerging evidence suggests that CD8a expression on NK cells may be associated with increased activation and improved receptor binding (21, 31). We noted increased degranulation of CD8^+^ NK cells (28.5%±15.7%; Figure 5B) This population further displayed increased expression (81.9%±8.6%) and MFI of CD16, as well as increased expression of CD161 (78.2%±14.7%). There was also an increased expression (96.7%±2.5) and MFI of CD45RA, but a decrease in the expression (2.5%±2.6%) and MFI of CCR7. The expression of the adenosine receptors CD39/CD73 was also decreased (8.3%±4.3%) in the CD8^+^ NK cell subset, consistent with a less tissue resident and more activated phenotype (40, 41).

**Figure 5.**
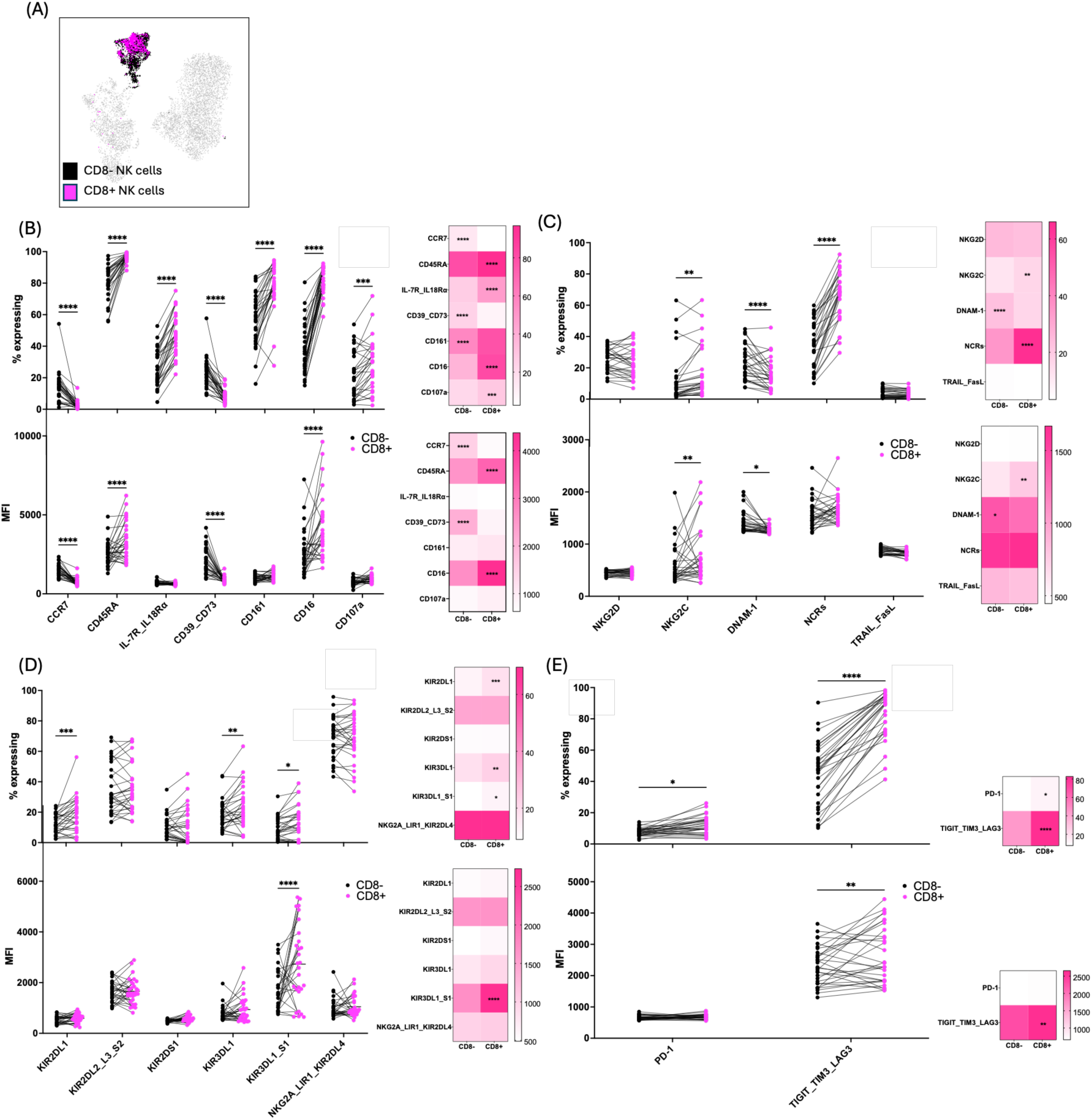
CD8^+^ NK cells have distinct phenotypes that reflect increased activation and maturation. A comparison of receptor expression between CD8- and CD8+ NK cell populations. (A) Overlay of CD8- and CD8+ populations on UMAP of the total lymphocyte population. (B-D) The percent expression (top graph) and MFI (bottom) of each receptor group. Lines connect same donors (B) Differential receptors. (C) Checkpoint receptors. (D) Activating receptors. (E) KIR expression. (n=30); statistics shown are a two-way ANOVA with multiple comparisons; *=p<0.05; **=p<0.01; ***= p < 0.001; ****=p<0.0001

The NCRs and NKG2C were the activating receptors more commonly expressed in the CD8^+^ population (NCRs: 66%±15.9%; NKG2C:16.3%±16.3%;) than the CD8^-^ NK cells (Figure 5C). In contrast, both the expression (16%±9.9%) and MFI of DNAM-1 were decreased on the CD8^+^ NK cells. There was no change in the expression of NKG2A/LIR1/KIR2DL4, but both KIR2DL1 (18.3%±11.4) and KIR3DL1 (23.1%±14%) had an elevated expression on CD8^+^ NK cells (Figure 5D). Both the expression of KIR3DL1_S1 (12.4%±11%) and the MFI was also elevated. Further, the expression of the checkpoint receptors was increased (PD-1: 12%±5.9; TIGIT/TIM3/LAG3: 82.6%±15.1%; Figure 5E), suggesting that these CD8^+^ NK cells have a more mature, activated phenotype.

### Educated NK cell responsiveness corresponds to the number of educating receptors on their cell surface

The education status of an NK cell can influence the level of responsiveness against a given target. Moreover, the strength of education can be impacted by the number of educating KIR on the NK cell surface (42). To assess how the expression of educating KIR impacted the responsiveness in our donor cohort, we compared degranulation of NK cells stimulated with K562 targets with increasing numbers of educating KIR on their cell surface (KIR2LD1, KIR2DL2/L3/S2, KIR3DL1, and/or NKG2A; Figure 6A). KIR negative NK cells had the lowest level of activation (9.2%±5.8%), with each of the subsequent groups displaying higher expression of CD107a (1 receptor: 22.9%±13.6%, p=0.07; 2 receptors: 31.9%±18.4%; 3 receptors:39.6%±20.3%; 4 receptors: 45.7%20.3%). The presence of two or more receptors trended toward greater responsiveness than having just one educating receptor on the NK cell surface (2 receptors: p=0.08; 3 receptors: p=0.0017; 4 receptors: p=0.07); this follows a known paradigm of NK cell education which demonstrates an additive impact on responsiveness with increasing expression of educated on the NK cell (43).

**Figure 6.**
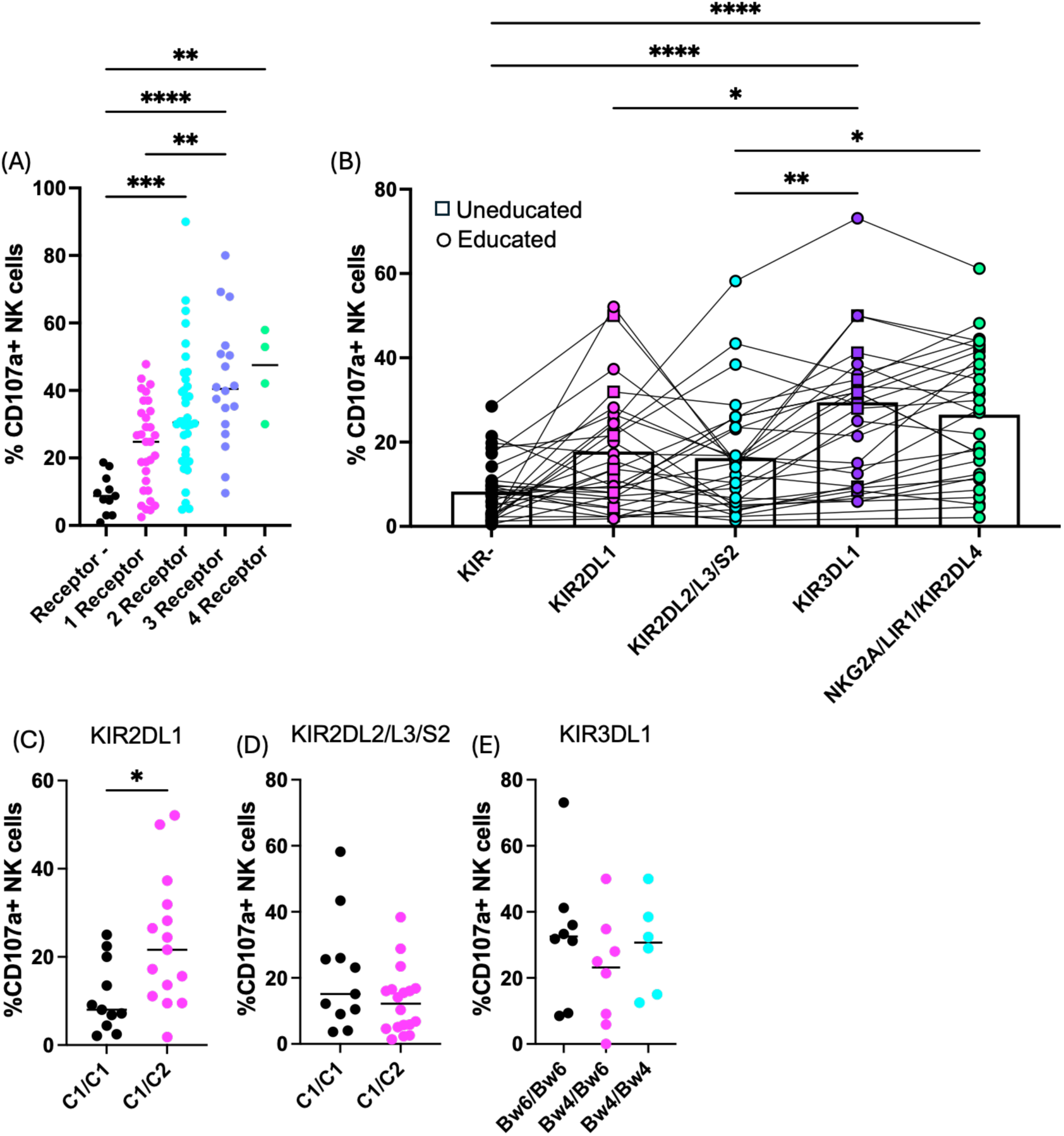
NK cell activation against an HLA-target increases as the number of educating KIR on a given NK cell increases. NK cell responsiveness was assessed by coculture with the HLA-K562 (3:1 E:T) and was measured by CD107a expression. (A) degranulation of educated NK cells relative to the number of educating receptors (KIR2DL1, KIR2DL2/L3/S2, KIR3DL1, NKG2A_LIR1_KIR2DL4) expressed by a given NK cell (n=11). Statistics shown are a one-way ANOVA with multiple comparisons. (B) degranulation of single positive KIR or NKG2A populations. Educated donors are denoted by a circle, uneducated denoted by a square (n=30). Lines connect same donors across KIR populations. Statistics shown are a one-way ANOVA with multiple comparisons. (C) degranulation of single positive KIR2DL1 in uneducated (C1/C1) or educated (C1/C2) subsets. Statistics shown are an unpaired t test. (D) degranulation of single positive KIR2DL2/L3/S2 in (C1/C1) or (C1/C2) subsets. Statistics shown are an unpair t test (E) degranulation of single positive KIR3DL1 in uneducated (Bw6/Bw6) or educated (Bw4/Bw6 or Bw4/Bw4) subsets. Statistics shown are a one-way ANOVA with multiple comparisons. *=p<0.05; **=p<0.01; ***= p < 0.001; ****=p<0.0001

To assess how any individual KIR influenced NK cell responsiveness, we examined CD107a expression on single positive KIR populations (Figure 6B). The single positive KIR3DL1^+^ (26.8%±17.9%) and NKG2A/LIR1/KIR2DL4+ (26.3%±15.7%) populations had comparable levels of responsiveness, and greater responsiveness than the KIR negative (8.2%±7%), KIR2DL1^+^ (17.6%±13.6%), and KIR2DL2/L2/S2^+^ populations (15.4%±13.4%). As education is determined by both the expression of a given KIR and it’s cognate HLA binding partner, we genotyped each donor to determine their expression of each of the KIR ligands (HLA subtypes with C1, C2 or Bw4 public epitopes then stratified them based on their education status.

KIR2DL1, which binds HLA-C2, had a greater responsiveness when the donor expressed HLA-C2 compared to donors who only expressed HLA-C1 (Figure 6C) KIR2DL2/L3/S2 primarily binds HLA-C1, though can weakly interact with HLA-C2; within our dataset, there was no difference in activation between donors that expressed HLA-C1 alone or with HLA-C2 (Figure 6D) (44, 45). Finally, KIR3DL1, educated by HLA-Bw4; we did not note changes in activation between donors that were or were not educated through KIR3DL1 (Figure 6E).

### T cell populations vary in their memory cell abundance

A key hallmark of both CD4^+^ and CD8^+^ T cells is their ability to form memory populations after antigen exposure. These memory subsets are phenotypically and functionally distinct. They are classically delineated by their expression of CCR7 and CD45RA into: naïve (CD45RA^+^CCR7^+^), central memory (CD45RA^-^CCR7^+^), effector memory (CD45RA^-^CCR7^-^), and terminally differentiated effector memory T cell re-expressing CD45RA (TEMRA; CD45RA^+^CCR7^-^)(17). Within our cohort, there was no difference between the CD4 and CD8 T cell naïve and effector memory populations (Figure 7A). There were a higher number of CD4^+^ central memory T cells (36.7%±10%) compared to the CD8^+^ central memory population (12.9%±7.7%), which reflects previous work showing a higher percentage of CD4^+^ memory T cells expressing CCR7(46, 47). In contrast, there were a higher number of CD8^+^ TEMRA cells (26.8%±11.7%) compared to the CD4^+^ TEMRA population (2.9%±1.9%). When dichotomized by age and sex, we noted no differences across the memory populations of CD4^+^ and CD8^+^ T cells (Figure 7B-E).

**Figure 7.**
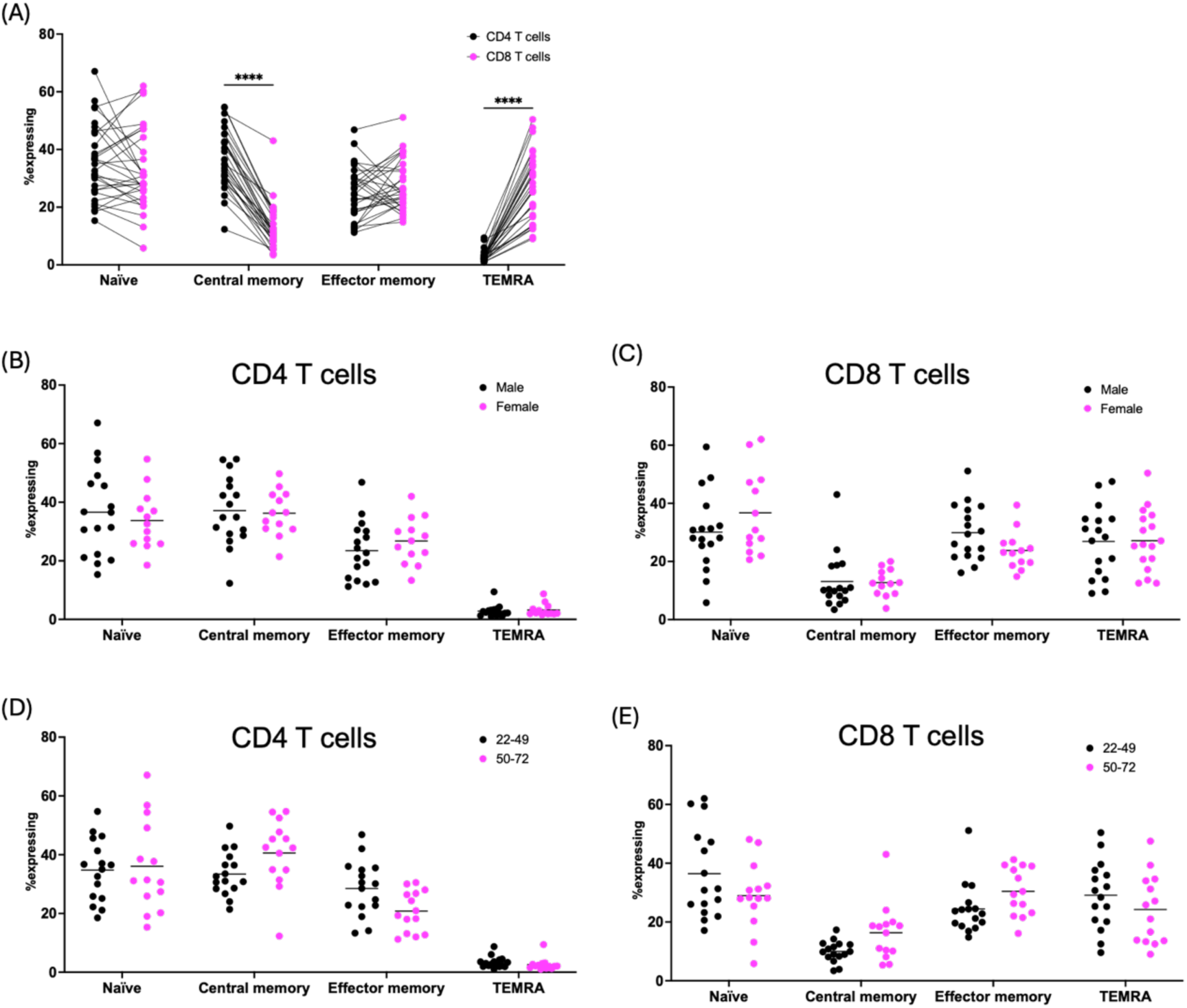
CD4^+^ and CD8^+^ T cells differ in their central memory and TEMRA memory cell compartments. (A) A comparison of the memory cell compartments between CD4 and CD8 T cells as determined by their expression of CD45RA and CCR7. Lines connect same donors. Statistics shown are a two-way ANOVA with multiple comparisons (B & C) Percent of each memory subset present dichotomized by sex ((B) CD4 T cells; (C) CD8 T cells) (D & E) Percent of each memory subset present dichotomized by age ((D) CD4 T cells; (E) CD8 T cells). (n=30); statistics shown are mixed-effects analysis with multiple comparisons; ****=p<0.0001

### CD8^+^ T cells acquire NK cell-like features with maturation

As the T cell memory repertoire undergoes differentiation and maturation, the receptor profile of each population can change. To determine how NK cell receptors fit with this paradigm, we next examined expression of receptors associated with NK cells and lymphocyte maturation among CD8^+^ T cell subsets. The naïve subset was notably distinct from the other memory subsets within the CD8^+^ T cell cluster on the UMAP (Figure 8A). There was some overlap between the central memory population and the effector memory population, but it was the effector memory and TEMRA populations that were most closely aligned. Expression of the adenosine receptors CD39/CD73 was highest in the naïve population (77.3%±9%) and steadily decreased as the T cell differentiated, with the lowest expression in the TEMRA population (29.8%±12%; Figure 8B) Reciprocally, the expression of CD161 was lowest among the naïve subset (2.6%±1.7%) and steadily increased with across the memory populations: central memory (9.1%±4.5%), effector memory (32.3%±13.1%), and TEMRA (34.6%±14.5), though effector memory subset displayed the highest MFI. This is in line with previous work showing CD161^+^ CD8^+^ T cells as a highly cytotoxic population(15). In agreement with the literature, the expression of the checkpoints was also elevated in the memory populations compared to naïve subsets– PD-1 was most highly expressed by the effector memory T cells (39.3%±11.3%), while TIGIT/TIM3/LAG3 was most highly expressed on the central memory population (50.5%±12.4%; Figure 8E)(48, 49).

**Figure 8.**
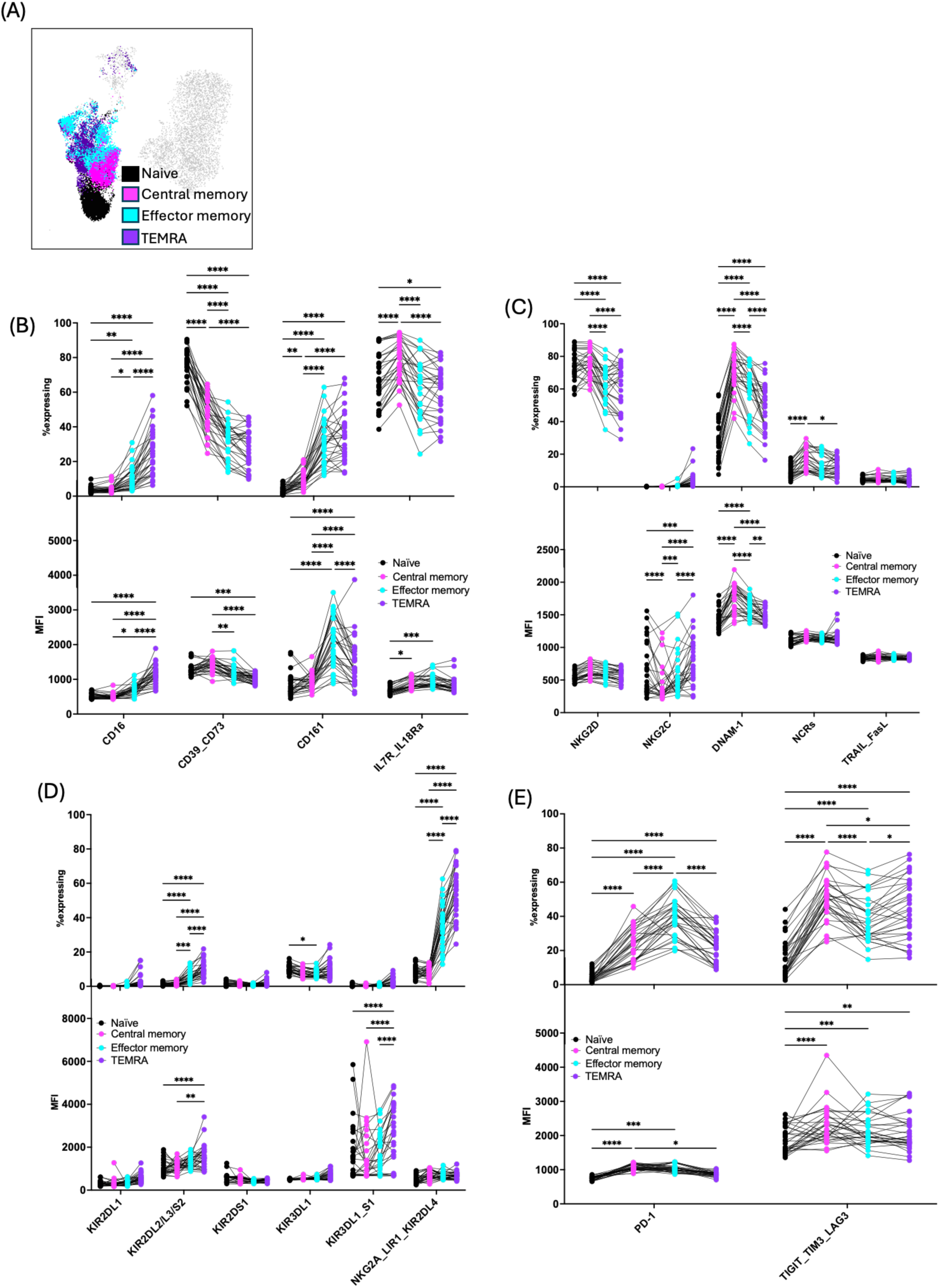
CD8^+^ T cells acquired NK cell-like features as they mature. A comparison of receptor expression within the memory cell compartments of CD8^+^ T cells determined by their expression of CD45RA and CCR7 (A) Overlay of the CD8^+^ T cell memory subsets on UMAP of the total lymphocyte population. (B-D) The percent expression (top graph) and MFI (bottom) of each receptor group. Lines connect same donors across memory populations (B) Differentiation receptors. (C) Activating receptors. (D) KIR expression. (D) Checkpoint receptors. (n=30); Statistics shown are two-way ANOVA with multiple comparisons; *=p<0.05; **=p<0.01; ***= p < 0.001; ****=p<0.0001

Further, within our donor cohort, there are several canonical NK cell receptors on CD8^+^ T cell memory populations. The expression of the Fc receptor CD16 was elevated in the effector memory (10.6%±6.8%) and TEMRA populations (25.2%±13.1%; Figure 8B). NKG2D was highly expressed across all the subsets, though naïve (74.1%±9.3%) and central memory T cells (76.3%±7.9%) had a greater percentage of NKG2D^+^ cells than effector memory (62.3%±10.4%) and TEMRA populations (60.8%±13.5%; Figure 8C). DNAM-1 expression was upregulated in the central memory subset (71.5%±11.1%) and more highly expressed in the effector memory (60.8%±14.4%) and TEMRA (49.9±14.3%) than the naïve T cells (27.8%±12.9). Moreover, expression of the NCRs peaked in the central memory population (16.2%±6.3%). On the side of inhibition, the expression of NKG2A_LIR1_KIR2DL4 was most highly expressed in the effector memory (34.7%±11.3%) and TEMRA populations (55.6%±14.1) compared to the naïve (7.6%±3.2%) and central memory populations (7.8%±3%; Figure 3.8 D). Finally, expression of KIR2DL1 was slightly upregulated in the TEMRA subset (3.2%±4.9%) and KIR2DL2/L3/S2 was elevated in the effector memory (5.3%±3.3%) and TEMRA populations (11.2%±6.9%).

Taken together, our findings indicated that there is some congruency in the receptors expressed by CD8^+^ T cells and NK cells as they reach a higher state of maturation and activation, indicating that cytotoxic lymphocytes likely share some conserved pathways of receptor acquisition that may improve their agility when responding to a challenge.

### CD4^+^ T cell effector memory subset has increased expression of DNAM-1 and the checkpoint receptors

To define putative use of NK cell receptors by CD4+ T cells during their differentiation, we assessed the expression of each receptor within the memory populations. The naïve subset once again formed a separate population from the other memory subsets within the CD4^+^ T cell cluster on the UMAP (Figure 9A). The central memory and effector memory populations had some overlap, with the former a potential transitional population between the naïve and effector cells, though this transition is not necessarily linear(50). TEMRA cells form a very small percentage of the CD4^+^ T cell population (3.3%±2.6%; Figure 7A) and are more diffuse throughout the UMAP (Figure 9A). In contrast to the CD8^+^ T cells, which lose CD39/CD73 with maturation, the CD4^+^ T cell maintain expression of these ATP-degrading/adenosine producing receptors across the subsets, with the effector memory population displaying the highest expression (46.9%±12.7; Figure 9B) (51, 52). However, like the CD8^+^ T cells, CD4^+^ T cells increase expression of CD161 with maturation, with the lowest expression in the naïve population (1.7%±0.9%), and a shift toward higher expression as they mature into the memory subsets, with highest expression found on the effector memory T cells (31.7%±8.6%). The CD4^+^ T cells further acquire expression of the checkpoint receptors as they mature, with the effector memory subsets expressing the highest percentage of both PD-1 (47.7%±9%) and TIGIT/TIM3/LAG3 (31.5%±9%; Figure 9E).

**Figure 9.**
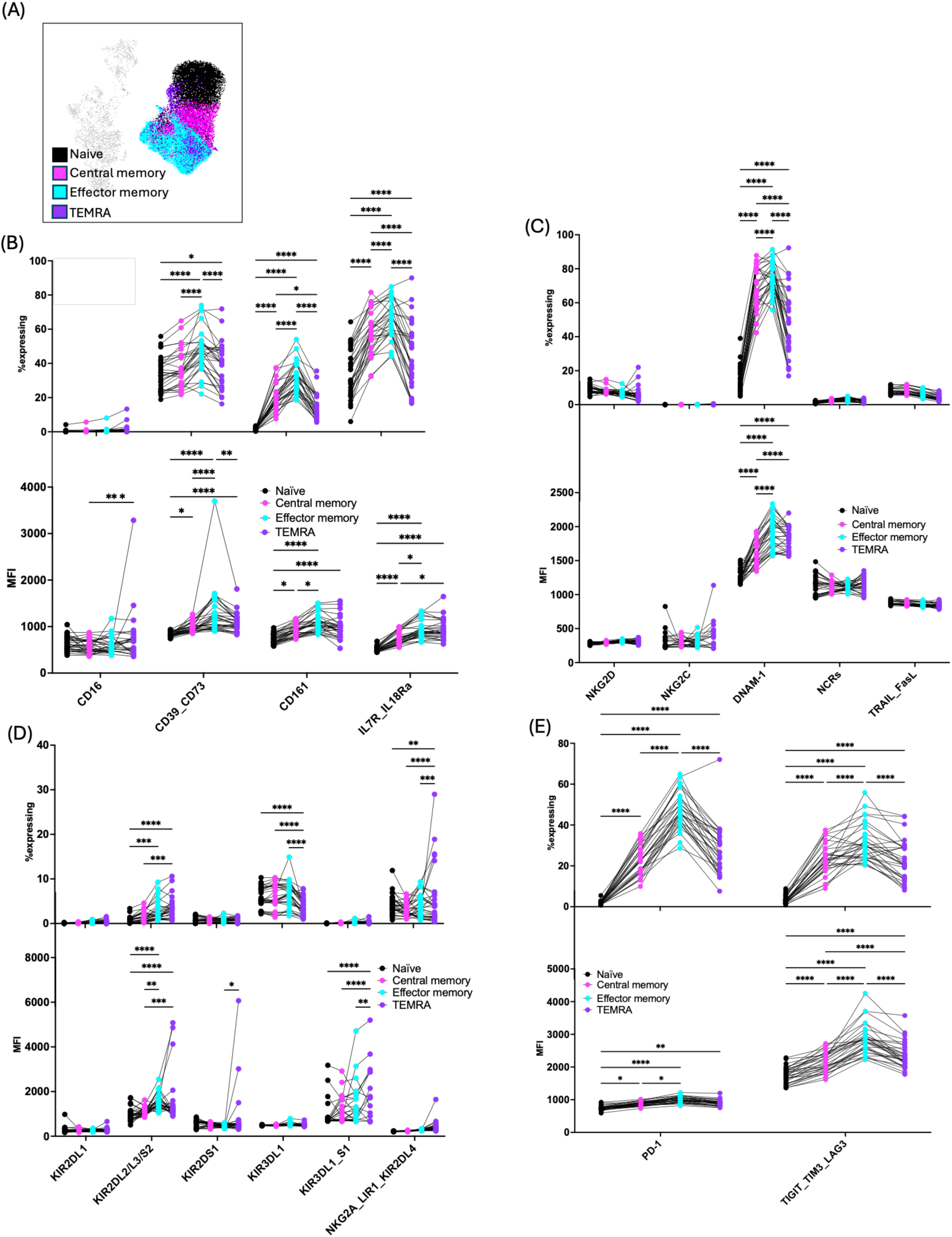
CD4^+^ T cells increase expression of checkpoint inhibitors as they mature. A comparison of receptor expression within the memory cell compartments of CD4^+^ T cells as determined by their expression of CD45RA and CCR7 (A) Overlay of the CD4^+^ T cell memory subsets on UMAP of the total lymphocyte population. (B-D) The percent expression (top graph) and MFI (bottom) of each receptor group. Lines connect same donors across memory populations (B) Differentiation receptors. (C) Activating receptors. (D) KIR expression. (E) Checkpoint receptors. (n=30); Statistics shown are two-way ANOVA with multiple comparisons; *=p<0.05; **=p<0.01; ***=p <0.001; ****=p<0.0001.

Among the activating receptors, only DNAM-1 changed between subsets, with the lowest expression in the naïve subset (15.2%±7.4%), and the central memory (68.4%±12.4%) and effector memory (75.1%±8.7%) subsets displaying the highest levels of expression. Noteworthy, the effector memory subset had the highest MFI, suggesting a higher overall density of this activating receptor (Figure 9C). Though KIR on CD4^+^ T cell is rarely reported, it has been found to be expressed on some CD4^+^ T cells populations are low levels.(53, 54) Within our cohort, we noted low level expression of KIR3DL1 across the subsets, with the highest expression in the central memory (6%±2.7%) and effector memory populations (6.5%±2.5%; Figure 9D). There was also low expression of KIR2DL2/L3/S2 in the effector memory (2.6%±2.2%) and TEMRA populations (3.4%±2.4%). Taken together, this data indicates that while CD4^+^ may express NK cell-associated inhibitory receptors at low levels, they do not share the same shift toward a more innate-like phenotype as effector memory and TEMRA CD8^+^ T cells.

## DISCUSSION

Though both arise from the common lymphoid progenitor, T and NK cells differ in their requirement for specific antigen and formation of memory subsets despite shared activation pathways and specific transcriptions factors as they mature (16). Associated with a more innate-like phenotype, NK cells are generally considered to be less antigen-restricted (often antigen-independent) counterpart to the specific antigen-dependent CD8^+^ T cells. We leveraged modern high-parameter flow cytometry techniques to examine the patterns of receptor expression in each of CD4^+^, CD8^+^ and NK subsets. For the first time, we create an atlas of receptor expression among these cell subtypes, focusing on canonically NK cell receptors. Here we have explored the expression patterns of 35 markers of lymphocyte differentiation, activation, and inhibition to better understand how these markers shape the T cell and NK cell repertoire in 25 FACS colours. We found that CD8^+^ T cells and NK cells share a breadth of activating and inhibitory receptors, that this receptor expression correlated with cell maturity, and that the patterns of both activating and inhibitory receptor expression were similar between the two cytotoxic lymphocyte subsets.

Within the NK cells, the expression of CD8^+^ was notably elevated on the CD56^dim^CD16^high^ population, in conjunction with an increase of KIR and CD161 expression. Further, the CD8^+^ NK cell population was associated with an increased degranulation against a K562 challenge. Within the CD8^+^ T cells, there was a high expression of NKG2D and DNAM-1. Further, both the effector memory and TEMRA CD8^+^ T cell populations displayed elevated expression of CD16, NKG2A/LIR1/KIR2DL4, and KIR2DL2/L3/S2, all markers notably ascribed as “NK cell receptors”. Unlike CD8^+^ T cells, CD4^+^ had limited expression of NK cell receptors, indicating that these expression patterns are shared by cytotoxic lymphocyte populations. Taken together, these data indicate that the receptor segregation classically applied to T cells and NK cells may not be as clearcut as previously thought and that there may be some congruency in how receptor expression changes as these cells mature and are activated.

CD8^+^ NK cells are associated with delayed HIV progression, reduced relapse risk in multiple sclerosis, and to correlate with patient outcomes in lymphoma (55–57). Several studies have demonstrated that CD8α homodimers can modulate NK cell activation (21, 31, 58, 59). CD8α expression is induced by IL-15 and marks a highly proliferative and more cytotoxic NK cell subset (21). Moreover, NK cells expanded *ex vivo* that express CD8α demonstrate improved serial killing (58). This may in part be because CD8-HLA I binding can reduce activation-induced apoptosis of NK cells, thus permitting their continued cytolytic functioning (59). On cytotoxic T cells, CD8 acts as a coreceptor that improves binding of the TCR to HLA I (60).

CD8α may be involved with KIR-HLA engagement on NK cells: CD8α is overexpressed on KIR^+^ NK cells and facilitates KIR-mediated inhibition, though it does not correlate with education (21). CD8α can act as a co-receptor for KIR3DL1 and improve binding to its ligand HLA-Bw4 (31). This correlation between CD8 and KIR expression was noted within our dataset, as CD8^+^ NK cells had higher expression of KIR2DL1 and KIR3DL1 populations.

The expression of NK cell-associated receptors has been previously noted on T cells, particularly CD8^+^ T cell subsets; these can modulate T cell activation. NKG2D and DNAM-1 can act as co-stimulators of effector CD8^+^ T cells, though not naïve T cells, and in some instances provide stronger co-stimulation than CD28 (61, 62). NKG2D is expressed by double negative thymocytes during T cell development and is required for the successful maturation of T cells into potent effector cells (63). It also influences the formation of memory effector T cells and can rescue memory recall responses in the absence of CD4^+^ T cell stimulation (22, 64).

Engagement of NKG2D on CD8^+^ T cells can trigger “bystander” activation, which enables a faster response against bacteria or viral pathogens because the ligands for NKG2D are upregulated by any cell in response to cellular stress (65). Indeed activation of CD8^+^ T cells via NKG2D co-stimulation has been noted in the context of infection, cancer, and autoimmunity (66–71). Noteworthy, most of the experimental work examining NKG2D^+^ T cells has been done with immunocompetent mouse models, and that the expression patterns of NKG2D vary between mouse and human T cells. Mouse NKG2D is only expressed after activation and not found on naïve subsets; in contrast, human CD8^+^ T cells constitutively express NKG2D (72). This study is the first comparison of NKG2D expression across healthy human CD8^+^ T cell memory populations. We note a decrease in the expression of NKG2D on effector memory and TEMRA CD8^+^ T cells. As this decrease in expression was contrasted by an increase in both checkpoint receptors and inhibitory NKG2A and KIR expression, more terminally differentiated T cells may shift to a less reactive phenotype, ostensibly to avoid autoimmunity or off-target effects (23, 73).

Immune checkpoints/inhibitory receptors seem to increase with increasing differentiation of lymphocytes. Among CD8^+^ T cells, PD-1, CTLA4 and others are co-expressed on antigen-specific activated T cells, suggesting a mechanism for their ultimate inhibition. Here, we find that NKG2A is also correlated with T cell maturation. Indeed, NKG2A has been reported as associated with persistent activation, T cell exhaustion, and decreased cytotoxicity (23, 74). The presence of NKG2A^+^ CD8^+^ T cells within solid tumours is associated with increased tumour progression and worse overall survival (75–77). Among NK cells, NKG2A is associated with less-mature (mainly cytokine-producing) subsets and it is KIR – a more powerful mediator of inhibition – that is associated with a more mature, and sometimes educated population; we observed this shift in expression with the CD56^dim^ CD16^high^. We find that, consistent with a hypothesis of increasing inhibitory receptor expression with maturation, TEMRA CD8^+^ T cell memory subsets that had higher expression of KIR2DL1 and KIR2DL2/L3/S2, along with higher expression of NKG2A. This co-expression of NKG2A and KIR on CD8^+^ T cells has previously been shown to identify a terminally differentiated, more-innate like subset with a higher expression of Eomesodermin and increased production of IFN-γ (78). However, another study of innate-like T cell found that NKG2A and KIR were expressed on functionally distinct subsets, with NKG2A^+^ T cells associating with increased IFN-γ production and CD161 expression, while KIR^+^ T cells were more cytotoxic and co-expressed CD57 and CD16 (24). Both CD16 and CD161 were upregulated in the effector memory and TEMRA subsets in our cohort, but we did not note the same segregation of markers (data not shown).

Though the acquisition of an NK cell-like phenotype is associated with T cell maturation and aging in general, of these receptors it was only DNAM-1 that was elevated on CD8^+^ T cells in the older cohort. In the context of age-related shifts in T cell population, there is conflicting evidence as to whether the naïve T cell compartment decreases with age; the population remains steady with CD4^+^ cells, but there has been some evidence of decrease with the CD8^+^ cells, though potentially only those infected with in cytomegalovirus (CMV) (79–82). We did not observe changes in the memory subsets between our younger and older cohorts, but it is important to note that our older cohort ranged from 50-72 years old, with only two donors aged 70 or older and only adults were included in our study. As we obtained our cohort from healthy individual donating blood to the Canadian Blood Services, we were limited by the demographics of those choosing and eligible to donate and did not have access to samples from people 75 or older. Many studies examining age-related changes in systemic T cell numbers considered older cohorts to be 65 or older and examine populations into their late 70’s and 80’s. This is similarly true for studies that have examined changes in the NK cell receptor repertoire as people age; we saw no changes in the receptor repertoire between the two age groups (83), and prevented us from studying the impacts of childhood on NK cell development, which are known to vary NK cell receptor usage (12).

This study leveraged high-parameter flow cytometry to examine the receptor repertoire of healthy human T cells, both CD8^+^ and CD4^+^, and NK cells from the peripheral blood to better understand the expression and co-expression patterns that shape a given cell’s function. We observed that mature NK cells expressed CD8^+^, had increased expression of KIR and checkpoint receptors, and are correlated with increased degranulation against an HLA I negative target cell. Moreover, we saw that CD8^+^ T cells acquired expression of NK cell associated receptors as they matured into a terminally differentiated phenotype, with increased expression of NKG2D, NKG2A, KIR, and CD16. As this study focused on the phenotypic changes within the immune cell subsets, we cannot draw functional conclusions for any given receptor. Nonetheless, the results of this work provide a gauge for the diversity and spread of receptor expression in a population of healthy donor PBMCs and can inform future work that examines the physiologic role of these receptors patterns.

## Supporting information

Lee Supplementary information

## ACKNOWLEDGEMENTS

This work was supported by an NSERC Discovery grant to JEB. SNL is a trainee of the Cancer Research Training Program of the Beatrice Hunter Cancer Research Institute. SNL is supported by a Canada Graduate Scholarship from the Canadian Institutes of Health Research and a Nova Scotia Graduate Scholarship through Dalhousie University. Dalhousie University is located in Mi’kma’ki, the unceded and ancestral territory of the Mi’kmaq people.

