## Supplementary material for "An atlas of natural killer cell receptor expression on healthy donor T and NK lymphocytes": Lee Supplementary information

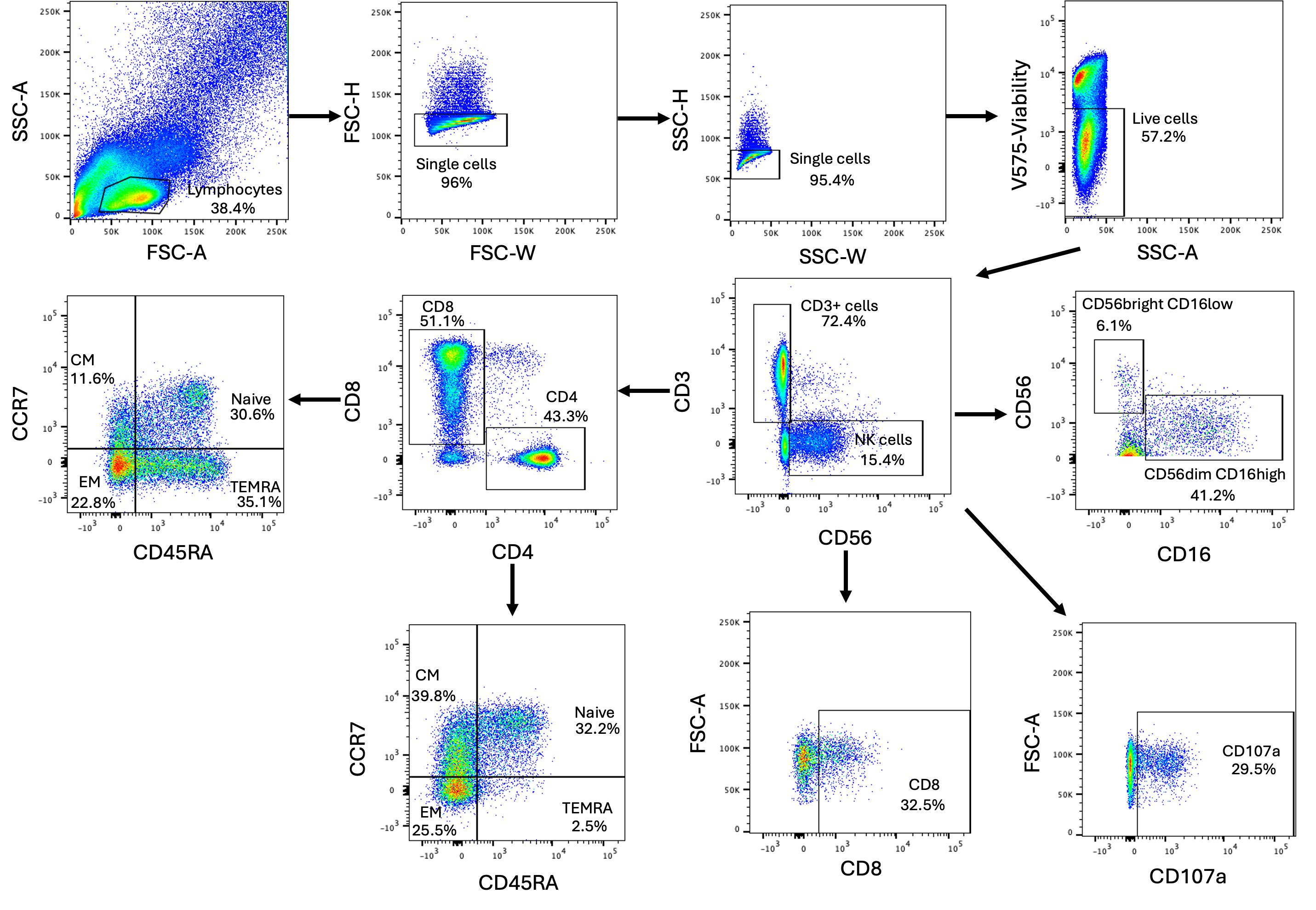

**Supplementary figure 1. Representative flow cytometry gating scheme for lymphocytes**

Lymphocytes were gated based on their size and complexity through the forward scatter area (FSC-A) and side scatter area (SSC-A), respectively. Duplicate cells were excluded by gating the single cells using the forward scatter height (FSC-H) and forward scatter width (FSC-W), followed by the side scatter height (SSC-H) and side scatter width (SSC-W). The cells were stained with the Viability Dye V575, which allowed for the exclusion of stained dead cells. NK cells were gated on as CD3^-^CD56^+^, followed by representative gating for CD56^bright^CD16^low^ vs CD56^dim^CD16^high^, CD107a, and CD8. T cells were gated on CD3^+^CD56^-^, followed by gating on CD8^+^ and CD4^+^. T cell memory populations were gated on CD45RA by CCR7

Supplementary table 1. Flow cytometry staining panel for human lymphocytes

| **Marker** | **Clone** | **Fluorochrome** | **Isotype** | **Company** | **Catalogue Number** |
| --- | --- | --- | --- | --- | --- |
| KIR3DL1 | DX9 | BV786 | Mouse | BD Biosciences | 742982 |
| CD39 | Tü66 | BV750 | Mouse | BD Biosciences | 747079 |
| CD73 | AD2 |  | Mouse | BD Biosciences | 747205 |
| TRAIL | RIK-2 | BV711 | Mouse | BD Biosciences | 743722 |
| FasL | NOK-1 |  | Mouse | BD Biosciences | 744101 |
| CD3 | UCHT-1 | BV650 | Mouse | BD Biosciences | 563852 |
| NKp46 | 9-E2 | BV605 | Mouse | BD Biosciences | 743710 |
| NKp30 | p30-15 |  | Mouse | BD Biosciences | 563384 |
| NKp44 | p44-8 |  | Mouse | BD Biosciences | 744301 |
| Live/Dead |  | FVS570 |  | BD Biosciences | 564995 |
| LAG-3 | T47-530 | BV480 | Mouse | BD Biosciences | 746609 |
| TIM-3 | 7D3 |  | Mouse | BD Biosciences | 746771 |
| TIGIT | 741182 |  | Mouse | BD Biosciences | 747843 |
| KIR2DS1 | 1127B | AF405 | Rabbit | R&D Systems | FAB8887V |
| NKG2C | 134591 | BB790-P | Mouse | BD Biosciences | Custom |
| NKG2D | 1D11 | BB660-P2 | Mouse | BD Biosciences | Custom |
| PD-1 | EH12.1 | BB630-P2 | Mouse | BD Biosciences | Custom |
| KIR3DL1/S1 | REA168 | FITC | REA | Miltenyi Biotech | 130-104-836 |
| KIR2DL1 | REA284 | PE-Cy7 | REA | Miltenyi Biotech | 130-120-447 |
| KIR2DL2/L3/S2 | GL183 | PE-Cy5.5 | Mouse | Beckman-Coulter | A66900 |
| CD161 | DX12 | PE-Cy5 | Mouse | BD Biosciences | 551138 |
| IL-7R | HIL-7R-M21 | PE | Mouse | BD Biosciences | 557938 |
| IL-18Ra | H44 |  | Mouse | BD Biosciences | 564675 |
| CD107a | H4A3 | APC-H7 | Mouse | BD Biosciences | 561343 |
| NKG2A | REA110 | APC | REA | Miltenyi Biotech | 130-113-563 |
| LIR-1 | GHI/75 |  | Mouse | Biolegend | 333720 |
| KIR2DL4 | REA768 |  | REA | Miltenyi Biotech | 130-112-466 |
| CD45RA | 5H9 | BUV805 | Mouse | BD Biosciences | 742052 |
| CD4 | SK3 | BUV737 | Mouse | BD Biosciences | 612748 |
| CCR7 | 2-L1-A | BUV661 | Mouse | BD Biosciences | 749824 |
| CD16 | 3G8 | BUV615 | Mouse | BD Biosciences | 751572 |
| CD8 | SK1 | BUV563 | Mouse | BD Biosciences | 741440 |
| DNAM-1 | DX11 | BUV496 | Mouse | BD Biosciences | 749935 |
| CD56 | NCAM16.2 | BUV395 | Mouse | BD Biosciences | 563554 |
